# Ferulic acid combined with skeletal stem cells to repair irradiated bone defect

**DOI:** 10.1101/2021.02.14.431131

**Authors:** Jia-Wu Liang, Pei-Lin Li, Qian Wang, Song Liao, Wei Hu, Zhi-Dong Zhao, Zhi-Ling Li, Bo-Feng Yin, Ning Mao, Li Ding, Heng Zhu

## Abstract

The reconstruction of irradiated bone defects after settlement of skeletal tumors remains a significant challenge in clinical applications. In this study, we explored radiation-induced skeletal stem cell (SSC) stemness impairments and rescuing effects of ferulic acid (FA) on SSCs *in vitro* and *in vivo*. The immunophenotype, cell renewal, cell proliferation, and differentiation of SSCs *in vitro* after irradiation were investigated. Mechanistically, the changes in tissue regeneration-associated gene expression and MAPK pathway activation in irradiated SSCs were evaluated. The regenerative capacity of SSCs in the presence of FA in an irradiated bone defect mouse model was also investigated. We found that irradiation reduced CD140a- and CD105-positive cells in skeletal tissues and mouse-derived SSCs. Additionally, irradiation suppressed cell proliferation, colony formation, and osteogenic differentiation of SSCs. The RNA-Seq results showed that tissue regeneration-associated gene expression decreased, and the western blotting results demonstrated the suppression of phosphorylated p38/MAPK and ERK/MAPK in irradiated SSCs. Notably, FA significantly rescued the radiation-induced impairment of SSCs by activating the p38/MAPK and ERK/MAPK pathways. Moreover, the results of imaging and pathological analyses demonstrated that FA enhanced the bone repair effects of SSCs in an irradiated bone defect mouse model substantially. Importantly, inhibition of the p38/MAPK and ERK/MAPK pathways in SSCs by specific chemical inhibitors partially abolished the promotive effect of FA on SSC-mediated bone regeneration. In summary, our findings reveal a novel function of FA in repairing irradiated bone defects by maintaining SSC stemness and suggest that the p38/MAPK and ERK/MAPK pathways contribute to SSC-mediated tissue regeneration post-radiation.

**Significance Statement:** Radiotherapy combined with surgery for the settlement of skeletal tumors usually leads to large bone defects and hampers wound healing. Skeletal stem cells (SSCs) have been defined as tissue-specific stem cells in skeletons and are highlighted in bone development and regeneration. Ferulic acid is a phytochemical with a wide range of therapeutic effects, including alleviation of radiation-induced oxidative stress and promotion of tissue regeneration. In the current study, promising data based on an *in vitro* cell model and an *in vivo* animal model demonstrates that ferulic acid alleviates radiation-induced impairment of SSCs and promotes SSC-mediated bone regeneration post-radiation.

## 1. Introduction

The settlement of skeletal tumors often requires therapeutic surgical bone removal in association with radiation therapy^1^. Although the strategy proved effective in the treatment of skeletal tumors, it usually results in large bone defects and hampered wound healing because of extensive tissue cutoff and irradiation-induced tissue damage. To aesthetically and functionally reconstruct bone defects and repair surrounding tissues, a number of techniques have been developed, including autogenous bone graft transplantation and microanastomosed free-flap reconstitution^2-5^. However, surgical reconstruction brings a high risk of morbidity due to prolonged anesthesia and secondary trauma.

In recent years, mesenchymal stem cell (MSC)-based tissue repair has been used for bone regeneration post-radiation but has yielded controversial results. Zuo et al. reported that rat bone marrow mesenchymal stem cell (BMMSC)-derived exosomes were capable of alleviating radiation-induced bone loss by restoring the function of recipient BMMSCs^6^. In another independent study, Liu et al. found that miR-34a promotes bone regeneration in irradiated bone defects by enhancing osteoblastic differentiation of BMMSCs in rats^7^. In contrast, Bléry et al. reported that the addition of a high concentration of BMMSCs did not improve bone regeneration in the irradiated areas of rats^8^. These controversial reports may result from unstandardized protocols in different labs. Notably, the role of native tissue-specific stem cells in skeletons, as the pivotal population contributing to bone formation and regeneration, was overlooked.

Skeletal stem cells (SSCs) have been recently characterized as skeletal tissue-specific stem cells that play pivotal roles in skeletogenesis and osteochondrogenic repair ^9-16^. Chan et al. identified a population of CD45-Ter119-Tie2-AlphaV+ multipotent stem cells in murine bone and revealed the presence of SSC niches that regulate their activity ^10^. A promising report from another lab demonstrated that Gremlin-1 defines a population of osteochondral reticular stem cells. Further functional studies suggest that these cells are closely involved in bone formation ^11^. Furthermore, the identification of self-renewing and multipotent SSCs in human bones validates the presence of tissue-specific stem cells in skeletal tissue and highlights their contribution to skeletal regeneration^16^. We have pursued the identification of tissue-specific stem cells from skeletal tissues and explored their application in the past decade. Our previous studies revealed the pivotal role of SSCs in controlling the inflammatory response by targeting host immune cells^17-23^. Additionally, our recent work demonstrated that SSCs play a pivotal role in the regulation of bone remodeling by suppressing inflammatory osteoclastogenesis via the concerted action of cell adhesion molecules and osteoprotegerin^24^. However, there is little information available about the change in SSC stemness, including self-renewal and multidifferentiation, after irradiation and the related underlying regulatory factors thus far, which limits the current level of understanding regarding SSC-based bone regeneration.

Ferulic acid (FA) is a widely distributed hydroxycinnamic acid that exhibits potent antioxidant and therapeutic activities^25^. FA has been widely applied in the prevention of reactive oxygen species-related diseases, including cardiovascular diseases, diabetes mellitus, and cancers^25^. In addition, FA has alleviated radiation-induced stem cell damage. Ma et al. performed hematopoietic progenitor colony-forming cell assays to assess the reconstitution of murine bone marrow after radiation-induced myelosuppression. They found that FA treatment enhanced hematopoietic progenitor cell activity and promoted blood cell recovery in mice after irradiation by cobalt-60 gamma resources^26^. Moreover, FA promoted *in vitro* osteogenic differentiation of BMMSCs by inhibiting micro340 to induce β-catenin expression through hypoxia^27^.

Given the fundamental role of SSCs in bone regeneration and the potential roles of FA in irradiation protection and osteogenic promotion, FA combined with SSCs may be an effective strategy for reconstructing irradiated bone defects. In the present study, we explore radiation-induced SSC impairments and the rescuing effects of FA on SSC-mediated bone regeneration by using an *in vitro* cell model and an *in vivo* animal model. Moreover, the cellular and molecular mechanisms underlying the protective effects of FA on SSCs were also investigated.

## 2. Materials and Methods

### Animals

Normal inbred 8-week-old C57BL/6 mice (n=90) and 2-week-old C57BL/6 mice (n=16) were purchased from Beijing Vital River Laboratory Animal Technology Co., Ltd. All of the experiments were performed in accordance with the Academy of Military Medical Sciences Guide for Laboratory Animals.

### SSC preparation

Murine SSCs were isolated from long bones according to our previously reported procedure^24^. Briefly, femurs and tibiae from 2-week-old C57BL/6 mice were dissected, and the bone marrow cells were flushed out before the long bones were chopped and digested by collagenase II (Sigma, 0.1% vol/vol) at 37 °C for 1 hour. The released cells were discarded, and the digested bone chips were cultured in minimum essential medium eagle, alpha modification (α-MEM; Invitrogen, Carlsbad, CA) supplemented with 10% fetal bovine serum (FBS) (HyClone, Logan, UT). The adherent cells from passages 3-6 were used for *in vivo* and *in vitro* experiments unless otherwise described. SSCs were also isolated from the femurs and tibiae of irradiated (cobalt-60) mice.

### FA preparation

Ferulic acid (98% purity, CAS 1135-24-6) was purchased from EFE BIO (Shanghai, China) and dissolved in DMSO to 20 mM as a stock solution. The optimized concentrations of FA were screened by Cell Counting Kit 8 (CCK-8) assay with graded concentrations of FA (0, 10, 20, 30 and 40 μM). It was determined that the optimal concentration of FA was 30μM. This concentration was used in the experiments unless otherwise specified. In some experiments, the SSCs were pretreated with FA for 24 hours before usage.

### Immunophenotyping of SSCs

To detect the effects of irradiation on the SSC immunophenotype *in vivo*, mice were irradiated with 2 Gy of gamma-radiation delivered at a rate of 0.98 Gy/minute. Femurs and tibiae were harvested 1 hour after irradiation for pathological analysis to determine the changes in CD140a-positive and CD105-positive cells in skeletal tissue. Additionally, SSCs were isolated from the long bones of irradiated mice (2 Gy) to investigate the SSC immunophenotype *in vitro*. For the mechanistic study, SSCs from normal mice were irradiated (2 Gy). Then, the SSCs were stained with PE- and PE-PerCP-conjugated monoclonal antibodies against mouse CD11b, CD31, CD44, CD45, CD80, CD105, CD140a and Sca-1 (eBio-Science, San Diego) according to the manufacturer’s protocol. Events were collected by flow cytometry with a FACSCalibur system (Becton-Dickinson), and data analysis was conducted with FlowJo 10.0 software.

### CCK-8 assay

The proliferation of SSCs was determined using the Cell Counting Kit 8 (CCK-8, Dojindo, Japan) according to the manufacturer’s protocol. Briefly, a total of 2×10^3^ irradiated SSCs or nonirradiated SSCs were seeded onto 96-well culture plates and added to the CCK-8 solution at a ratio of 100 μL:1 μL, and the plates were incubated at 37 °C for 1 hour. Absorbance was then measured at a wavelength of 450 nm using a microplate reader. Five wells at each time point were assayed. The CCK-8 experiments were performed at different timepoints. FA was added at graded concentrations (10, 20, 30 and 40 μM) to investigate the potential effects of FA on SSC proliferation.

### Colony-forming unit fibroblast formation assay (CFU-F assay)

In the current study, SSC self-renewal was assessed by a CFU-F assay. Aliquots (2×10^2^/well) of irradiated SSC or nonirradiated SSC suspensions were added in triplicate into six-well culture plates and maintained in culture for 14 days. After crystal violet staining, visible colonies that were larger than 3 mm in diameter were counted using an inverted microscope. FA was added at graded concentrations to investigate the potential effects of FA on the self-renewal of SSCs.

### Multilineage differentiation experiments

The osteogenic and adipogenic differentiation of irradiated SSCs and nonirradiated SSCs was determined by induction agents as previously described (10,16). For osteogenic differentiation, SSCs were seeded into 48-well plates at a cell density of 2×10^3^/well (five wells in each group) and incubated in osteogenic induction medium (10 mM glycerol-2-phosphate, 0.1 mM dexamethasone, and 20 mM ascorbic acid) for 14 or 28 days. For adipogenic differentiation, SSCs were seeded into 48-well plates at a cell density of 1×10^4^/well (five wells in each group) and incubated in adipogenic induction medium for 14 days (1 mM isobutylmethylxanthine and 10-3 mM dexamethasone). To evaluate osteogenesis at day 14, the expression of the osteogenic marker alkaline phosphatase (ALP) was assessed using a histochemical kit (Sigma) per manufacturer’s instructions. To evaluate adipogenesis at day 14, Oil Red O staining was performed according to previously described methods. To further determine the multiple-differentiation capacity of SSCs, the mRNA expression levels of the osteogenic genes Runx-2 and OCN and the adipogenic genes CEBP/α and PPARγ were determined at day 10 using quantitative polymerase chain reaction (qPCR). FA was added at graded concentrations to investigate the potential effects of FA on the multidifferentiation of SSCs.

### RNA sequencing analyses of SSCs

To further investigate the underlying mechanism of irradiation-mediated impairment of SSC properties, SSCs from irradiated or nonirradiated mice (2 Gy, 1 hour after irradiation) were isolated, and high-throughput RNA sequencing analyses were performed. Next-generation sequencing library preparations and Illumina MiSeq sequencing were conducted at GENEWIZ, Inc. (Suzhou, China). DNA libraries were validated by an Agilent 2100 Bioanalyzer (Agilent Technologies, Palo Alto, CA, USA) and quantified with a Qubit 2.0 Fluorometer. DNA libraries were multiplexed and loaded onto an Illumina MiSeq instrument per manufacturer’s instructions (Illumina, San Diego, CA, USA). Sequencing was performed using a 2×300 paired-end (PE) configuration; image analysis and base calling were conducted by MiSeq Control Software (MCS) embedded in the MiSeq instrument. All differentially abundant mRNAs were used for GO analysis to deepen the understanding of the molecular mechanism of cell biological information processes.

### SSC-microcryogel preparation

To facilitate SSC transplantation, SSC microcryogels were prepared according to previous reports^28,29^. Biodegradable gelatin microcryogels were obtained from Beijing Cytoniche Co., Ltd. (http://www.cytoniche.com/). Briefly, irradiated SSCs or FA-pretreated irradiated SSCs were harvested and resuspended at a concentration of 1×10^7^ cells/ml. From each of these suspensions,100-μL was pipetted onto a dispersible and dissolvable porous microcarrier tablet (3D Table Trix, 20 mg/tablet) to allow the suspension to be directly absorbed and hydrate the porous structures. The SSC microcryogels were then maintained in a humidified chamber and incubated at 37 °C for 2 hours to allow for further cell attachment. After that, PBS was added to develop an injectable SSC microcryogel at a concentration of 20 mg microcarrier/ml PBS.

### Transplantation of the SSC microcryogel in the irradiated bone defect murine model

To explore the repairing capacity of SSC on irradiated bone defects, the femurs of mice were irradiated (2 Gy). Bone defect surgeries were conducted 1 hour after irradiation. A 1.5-mm femoral defect was generated in both femurs. Nonirradiated mice that underwent surgery served as bone defect controls. A 50-μL SSC microcryogel was injected into the local bone defect of irradiated mice unless otherwise described. The same concentration of microcryogel without SSCs was used as a control. To study the effects of FA on bone repair, FA solution was injected into irradiated bone defects. Mice were sacrificed at 1, 2, and 3 weeks post-surgery, and the femurs were harvested for further micro-CT analysis and pathological evaluation.

### µ computerized tomography (μCT) analysis

The murine femurs were harvested and fixed overnight in 10% neutral buffered formalin, washed twice in PBS, and kept in 70% ethanol at 4°C. μCT scanning and analysis were performed using a Scanco μCT-40 (Scanco Medical). The femurs were scanned at a resolution of 8 μm, and reconstruction of three-dimensional (3D) images was performed using a standard convolution back-projection. The bone volume/tissue volume ratio (BV/TV) was calculated by measuring 3D distances directly in the trabecular network.

### Histological Examination and Immunohistochemistry

All femurs were fixed in 4% paraformaldehyde for 3 days, decalcified in 10% EDTA solution for 7 days, and then embedded in paraffin. The femurs were sectioned into 6-mm slices that were subjected to HE and Masson staining. In addition, osteogenic regeneration and SSC involvement were further evaluated by immunohistochemical analyses of Col-I and OCN.

### Real-Time Quantitative Polymerase Chain Reaction

qPCR was performed to further investigate SSC differentiation and its underlying mechanisms. The procedure was performed according to previous studies. Briefly, total RNA was extracted using TRIzol reagent (Fermentas) and reverse transcribed using the mRNA Selective PCR Kit (TaKaRa) according to the manufacturer’s instructions. Murine Runx2, OCN, CEBP/α, PPARγ, Fgf2, Stc1, Bmpr1b, Clec3b, Lama5, Gata2, Nanog, and Sox2 cDNA were amplified by real-time qPCR using a SYBR PCR Master Mix Kit (Sigma-Aldrich) and a 7500 Real-Time PCR Detection System (Applied Biosystems, ABI). The primers were synthesized by Invitrogen and are shown in supplementary Table 1. The mRNA levels were normalized to the value of glyceraldehyde-3-phosphate dehydrogenase.

### Western-blotting

Normal mouse-derived SSCs were seeded into cell culture plates (5×10^5^ cells/well) and irradiated with 2 Gy Co 60. FA was added to the SSC culture at the earliest possible time after irradiation. Nonirradiated SSCs were used as control. The SSCs were collected at 60 min post-irradiation. Protein lysis buffer (BioRad, Hercules, CA) was added, and the thawed lysates were vortexed and centrifuged. Proteins were separated by 10% sodium dodecyl sulfate-polyacrylamide gel electrophoresis (SDS-PAGE) and transferred onto nitrocellulose membranes. The membranes were blocked by incubation with 5% wt/vol nonfat dry milk. The membranes were then incubated with anti-phospho-JNK (P-JNK), anti-JNK, anti-phospho-p38 (P-p38), anti-p38, anti-phospho-ERK (P-ERK), anti-ERK, and anti-GAPDH antibodies (Cell Signaling Technology) at the appropriate dilutions overnight at 4°C. After incubation, the membranes were washed in Tris buffered saline with Tween-20 (TBST). Horseradish peroxidase-conjugated secondary antibodies were added to the membranes in 5% nonfat dry milk in TBST.

### Statistical analysis

Data are represented as the mean values with standard deviations. Statistical significance was analyzed using Student’s t-test. P values less than 0.05 were considered significant.

## 3. Results

### Irradiation-induced stemness impairment of skeletal stem cells

CD105 and CD140a are pivotal cell surface markers to identify tissue specific stem/progenitor cells in skeletons as previously reported^18,19,23,24^. As shown in Figure 1A and 1B, 2 Gy irradiation caused a significant reduction in the number of *in situ* CD105^+^ and CD140a^+^ cells by > 30% in murine femurs (\**p* < 0.05, ** *p* < 0.01, *** *p* < 0.001). Additionally, the results of immunohistochemical staining of bone slices demonstrated that irradiation led to a decrease in the number of CD105^+^ and CD140a^+^ cells in murine femurs in a radiation dose-dependent manner (Figure 1A and 1B). To minimize secondary effects *in vivo*, murine SSCs were isolated 1 hour post-irradiation. The immunophenotype, colony formation, cell proliferation, and osteo-adipogenic differentiation were evaluated. The results of immunophenotyping showed that irradiation (2 Gy) remarkably decreased the ratios of CD105 and CD140a cells in SSCs (Figure 1C) (** *p* < 0.01). In addition, the CFU-F assay showed that irradiated SSCs exhibited decreased colony formation capacity relative to that of nonirradiated SSCs (Figure 1D and 1E) (\**p* < 0.05, ** *p* < 0.01, *** *p* < 0.001). Consistent with the results of CFU-F experiments, the expression of self-renewal-related genes, including Nanog and Sox2, remarkably decreased after irradiation (Figure 1F)(** *p* < 0.01, *** *p* < 0.001). Furthermore, the CCK-8 data demonstrated that irradiation inhibited cell proliferation (Figure 1G) (** *p* < 0.01, *** *p* < 0.001). Moreover, irradiation inhibited ALP activity (Figure 1H) and the expression of osteogenic genes, including Runx-2 and OPN (Figure 1I), in SSCs. The suppressive effect was dose-dependent upon irradiation (Figure 1H and 1I)(\**p* < 0.05, ** *p* < 0.01, *** *p* < 0.001). No obvious influences of irradiation (2Gy)on adipogenic differentiation of SSC were observed (Figure S1A and S1B). Therefore, these results suggest that irradiation caused significant impairment of SSC stemness.

**Figure 1.**
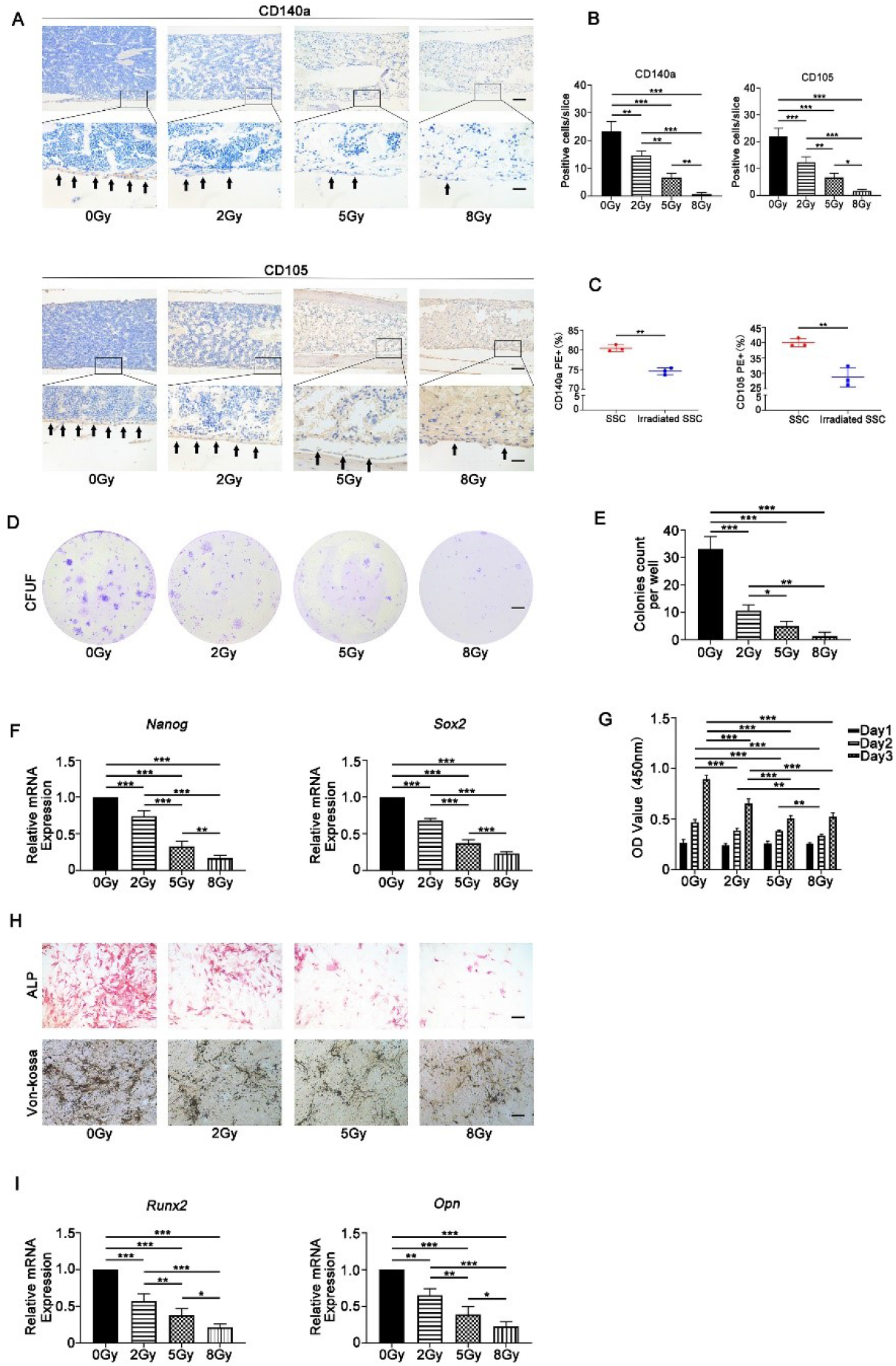
Irradiation caused stemness impairment of skeletal stem cells. Irradiation caused a significant reduction in the number of *in situ* CD105^+^ and CD140a^+^ cells in murine femurs at a radiation dose-dependent manner (Figure 1A and 1B). Additionally, irradiation remarkably decreased the ratios of CD105 and CD140a cells in SSCs *in vitro* (Figure 1C). Furthermore, the CFU-F assay showed that irradiated SSCs exhibited decreased colony formation capacity relative to that of nonirradiated SSCs (Figure 1D and 1E). Moreover, the expression of self-renewal-related genes, Nanog and Sox2, remarkably decreased after irradiation (Figure 1F). The CCK-8 data demonstrated that irradiation inhibited cell proliferation (Figure 1G). Further data showed that irradiation inhibited ALP activity (Figure 1H) and the expression of osteogenic genes, including Runx-2 and OPN (Figure 1I), in SSCs at dose-dependent manner upon irradiation (Figure 1H and 1I). Bars in Figure 1A, 1D and 1H represent 200μm, 500μm and 200μ, respectively. \**p* < 0.05, ** *p* < 0.01, *** *p* < 0.001.

### Differential gene expression and activation of MAPK pathways characterize the response of skeletal stem cells to irradiation

To explore the underlying mechanisms by which irradiation impaired SSC properties, the mRNA expression profile of SSCs after irradiation was identified by transcriptome sequencing. A total of 55,574 genes were examined in the SSC (n = 3) group and irradiated SSC (2 Gy, n = 3) group. The false discovery ratio was less than 0.05. The difference multiple was greater than 2-fold change to define significantly different proteins. A total of 638 genes were upregulated and 198 genes were downregulated in the irradiated SSC group compared with the SSC control group. The data in Figure S2A and S2B present the differentially expressed genes in heatmap and volcano plot format between SSC and irradiated SSC. Notably, the expression of osteogenesis-related genes, including Fgf2, Stc1, Bmpr1b, and Clec3b, and the cell proliferation-related genes Lama5 and Gata2 were significantly downregulated (Figure 2A). The results of Gene Ontology (GO) analysis showed that all of the differentially abundant genes between the SSCs and the irradiated SSCs were mainly involved in ossification, regulation of ossification, osteoblast differentiation, biomineral tissue development, and bone mineralization, etc (Figure 2B). The results of qPCR validated the gene expression of RNA sequencing (Figure 2C)(** *p* < 0.01, *** *p* < 0.001).

**Figure 2.**
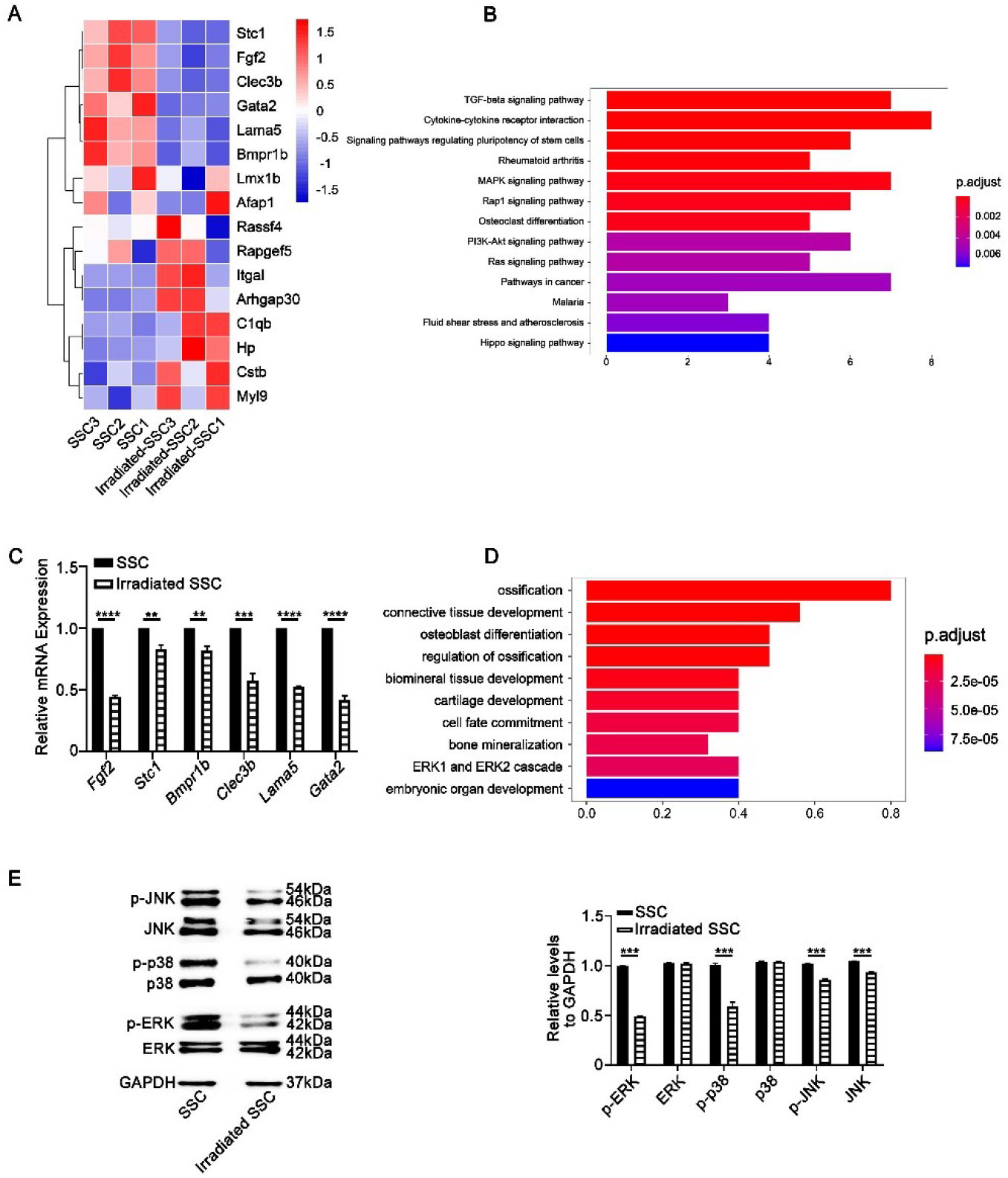
Differential gene expression and activation of MAPK pathways characterize the response of skeletal stem cells to irradiation. The results of mRNA sequencing (2Gy, n=3) showed that the expression of osteogenesis-related genes, including Fgf2, Stc1, Bmpr1b, and Clec3b, and the cell proliferation-related genes Lama5 and Gata2 were significantly downregulated (Figure 2A). The results of Gene Ontology (GO) analysis showed that all of the differentially abundant genes between the SSCs and the irradiated SSCs were mainly involved in ossification, regulation of ossification, osteoblast differentiation, biomineral tissue development, and bone mineralization, etc (Figure 2B). The data of qPCR validated the gene expression of RNA sequencing (Figure 2C). Further KEGG analysis suggested that the MAPK signaling pathway, TGF-β signaling pathway and cytokine-cytokine receptor signaling pathway were closely involved in the irradiation-induced changes in SSC properties (Figure 2D). The western-blotting data showed that irradiation (2 Gy) significantly inhibited the phosphorylation of p38/MAPK and ERK/MAPK in SSCs. There was obvious suppression of both the phosphorylated JNK and total JNK proteins in irradiated SSCs (Figure 2E) ** *p* < 0.01, *** *p* < 0.001.

Further KEGG analysis suggested that the MAPK signaling pathway, TGF-β signaling pathway and cytokine-cytokine receptor signaling pathway were closely involved in the irradiation-induced changes in SSC properties (Figure 2D).

Based on the KEGG analysis results and the important roles of MAPK signaling in bone formation and response to stress, the phosphorylation of the p38/MAPK, ERK/MAPK, and JNK/SAPK pathways was investigated. The results showed that irradiation (2 Gy) significantly inhibited the phosphorylation of p38/MAPK and ERK/MAPK in SSCs. In addition, there was obvious suppression of both the phosphorylated JNK and total JNK proteins in irradiated SSCs (Figure 2E) (*** *p* < 0.001).

### Ferulic acid (FA) partially alleviates irradiation-induced stemness impairment of skeletal stem cells

Because FA has been proven to be closely involved in radiation protection and bone formation in previous studies^26,27^, we investigated its role in the regulation of SSC stemness in the current study. As shown in Figure 3A, FA promoted SSC proliferation in an FA dose-dependent manner (\**p* < 0.05, ** *p* < 0.01, *** *p* < 0.001). Notably, the CCK-8 results demonstrated that FA significantly rescued the irradiation-induced proliferation inhibition of SSCs (Figure 3A) (\**p* < 0.05, ** *p* < 0.01, *** *p* < 0.001). Furthermore, ALP activity assays showed that FA (30 μM) promoted osteogenic differentiation of SSCs and irradiated SSCs (Figure 3B). The real-time PCR results demonstrated that FA enhanced the expression of osteogenic Runx-2 and OPN in SSCs (Figure 3C) (** *p* < 0.01, *** *p* < 0.001). The colony formation capacity partially reflects SSC stemness. We found that in the presence of FA, SSCs developed more CFU-F than their counterparts in the absence of FA (Figure 3D) (*** *p* < 0.001). Importantly, FA addition partially restored the colony formation that was compromised by irradiation, indicating that FA alleviates irradiation-induced SSC stemness impairment (Figure 3D). Moreover, FA regulated the expression of the self-renewal-related genes Nanog and Sox2 in SSCs and irradiated SSCs (Figure 3E) (\**p* < 0.05, ** *p* < 0.01, *** *p* < 0.001).

**Figure 3.**
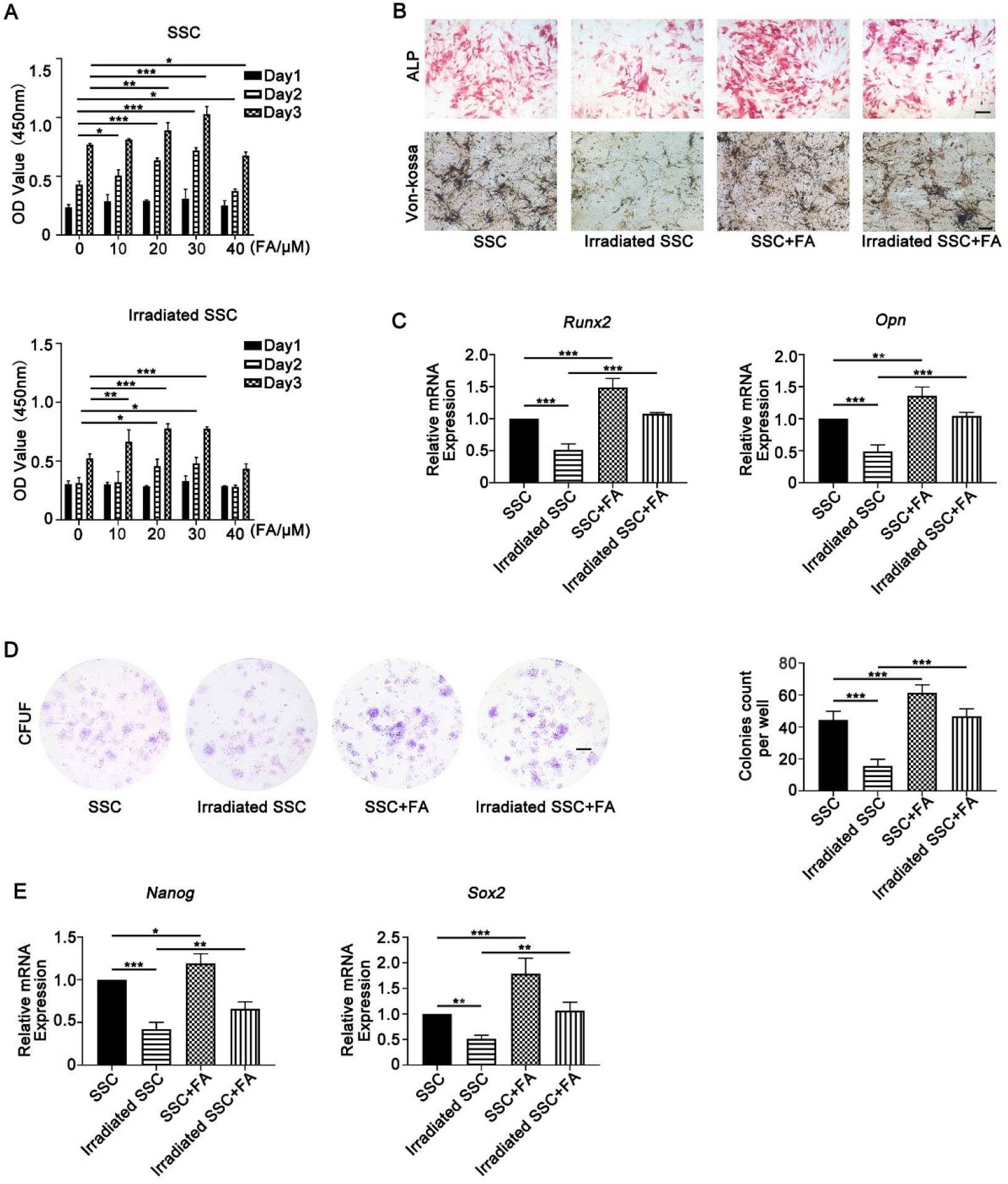
Ferulic acid (FA) partially alleviates irradiation-induced stemness impairment of skeletal stem cells. FA promoted SSC proliferation in an FA dose-dependent manner (Figure 3A). Notably, the CCK-8 results showed that FA significantly rescued the irradiation-induced proliferation inhibition of SSCs (Figure 3A). Furthermore, the data of ALP staining showed that FA (30 μM) promoted osteogenic differentiation of SSCs and irradiated SSCs (Figure 3B). The qPCR results demonstrated that FA enhanced the expression of osteogenic Runx-2 and OPN in SSCs (Figure 3C). Moreover, SSCs developed more CFU-F in the presence of FA (Figure 3D). Importantly, FA supplementation partially restored the colony formation that was compromised by irradiation (Figure 3D). Moreover, FA regulated the expression of the self-renewal-related genes Nanog and Sox2 in SSCs and irradiated SSCs (Figure 3E). Bars in Figure 3B and 3D represent 200μm and 500μm, respectively. \**p* < 0.05, ** *p* < 0.01, *** *p* < 0.001.

### FA partially rescues irradiation-induced stemness impairment of SSCs by activating the p38/MAPK and ERK/MAPK pathways

To explore the association of the MAPK pathways and the promising FA-mediated effects on SSC, FAs were added to the SSC culture system. As shown in Figure 4A, supplementation with FA (30 μM) significantly restored the phosphorylation of p38/MAPK and ERK/MAPK in irradiated SSCs (Figure 4A)(\**p* < 0.05, *** *p* < 0.001). However, no obvious effects of FA on the phosphorylated JNK and total JNK proteins in nonirradiated SSCs and irradiated SSCs were observed.

**Figure 4.**
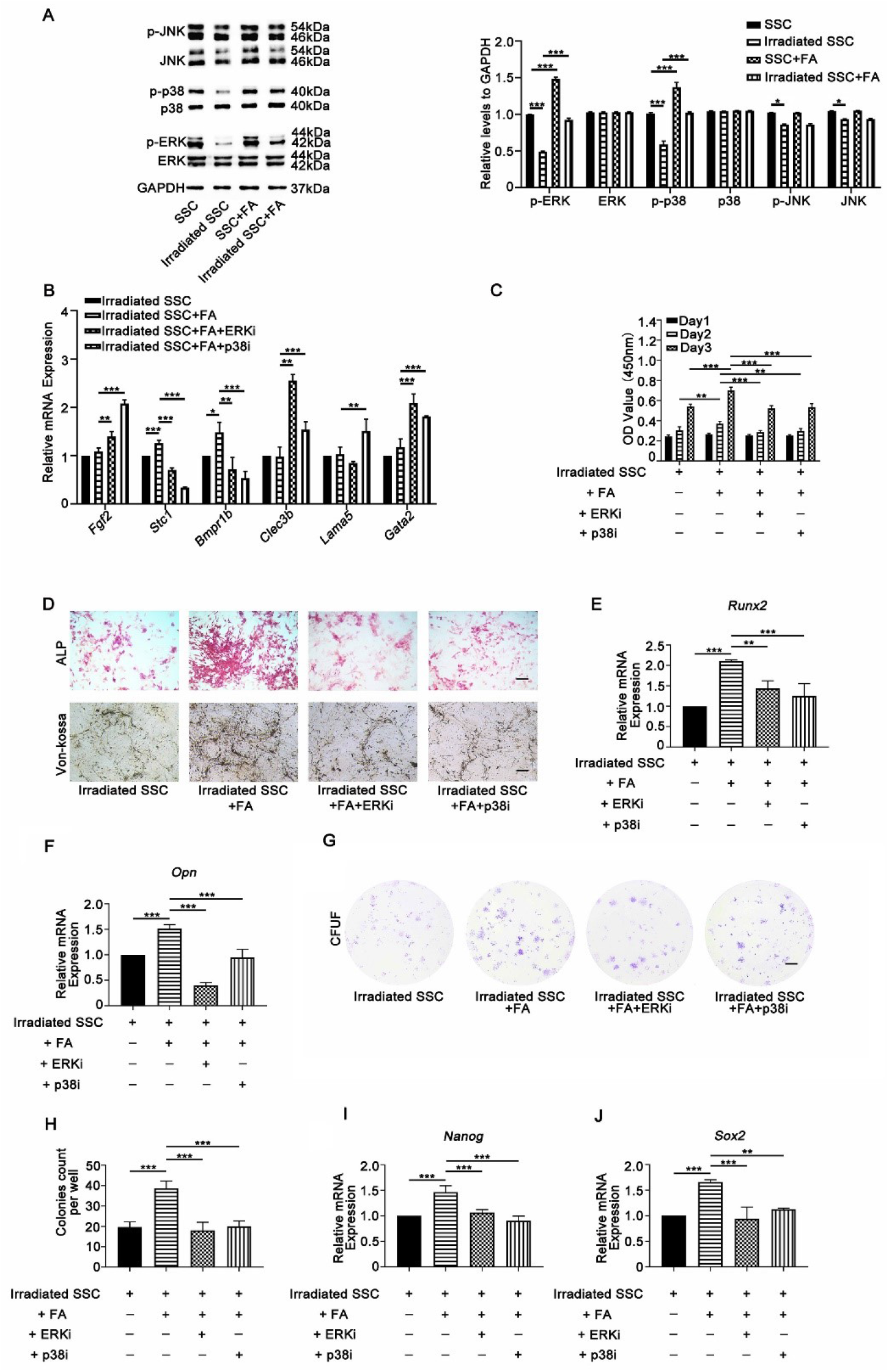
FA partially rescues irradiation-induced stemness impairment of SSCs by activating the p38/MAPK and ERK/MAPK pathways. Supplementation with FA (30 μM) significantly restored the phosphorylation of p38/MAPK and ERK/MAPK in irradiated SSCs (Figure 4A). No obvious effects of FA on the phosphorylated JNK and total JNK proteins in nonirradiated SSCs and irradiated SSCs were observed. Inhibition of p38/MAPK and ERK/MAPK signaling by specific inhibitors (SB203580 and PD98059) abolished the promoting effects of FA on the mRNA expression of Stc1 and Bmpr1b. Additionally, the CCK-8 assay data demonstrated that SB203580 and PD98059 blocked the rescued effects on cell proliferation in irradiated SSCs (Figure 4C). Moreover, SB203580 and PD98059 significantly suppressed ALP activity and the mRNA expression of Run2 and OPN in irradiated cells in the presence of FA (Figure 4D, 4E and 4F). Further, the colony numbers of irradiated SSCs in the FA-buffered culture system, and the mRNA expression of Nanog and Sox2 dramatically decreased in the presence of SB203580 and PD98059, respectively (Figure 4G, 4H, 4I and 4J). Bars in Figure 4D and 4G represent 200μm and 500μm, respectively. \**p* < 0.05, ** *p* < 0.01, *** *p* < 0.001.

To further clarify the innate relations of FA-induced activation of MAPK signaling and SSC biological properties, specific chemical inhibitors, including SB203580 (for p38/MAPK) and PD98059 (for ERK/MAPK), were added to the SSC culture system. The expression of genes that control SSC proliferation and differentiation, cell proliferation, osteogenic differentiation, and the colony formation capacity of SSCs were evaluated. The real-time results in Figure 4B show that inhibition of p38/MAPK and ERK/MAPK signaling by specific inhibitors abolished the promoting effects of FA on the mRNA expression of Stc1 and Bmpr1b, indicating that these genes may be involved in FA-mediated regulation of SSC osteogenic differentiation and cell proliferation via MAPK signaling (\**p* < 0.05, ** *p* < 0.01, *** *p* < 0.001). Additionally, the CCK-8 assay data demonstrated that SB203580 and PD98059 blocked the rescued effects on cell proliferation in irradiated SSCs substantially (Figure 4C) (** *p* < 0.01, *** *p* < 0.001). Moreover, SB203580 and PD98059 significantly suppressed ALP activity and the mRNA expression of Run2 and OPN in irradiated cells in the presence of FA (Figure 4D, 4E and 4F) (** *p* < 0.01, *** *p* < 0.001). Further, the colony numbers of irradiated SSCs in the FA-buffered culture system dramatically decreased in the presence of SB203580 and PD98059 (Figure 4G and 4H)(*** *p* < 0.001). Consistent with the change in colony formation assays, the mRNA expression of Nanog and Sox2 was also downregulated with the addition of specific inhibitors of the p38/MAPK and ERK/MAPK pathways, respectively (Figure 4I and 4J) (** *p* < 0.01, *** *p* < 0.001). Thus, these data suggest that FA enhanced osteogenic differentiation and proliferation partially via activation of the p38/MAPK and ERK/MAPK pathways.

### FA primed SSCs combined with microcryogel promotes bone repair in a murine irradiated bone defect model

Although *in vitro* studies have shown that FA is capable of maintaining SSC stemness, it remains unknown whether this is the case *in vivo*. Therefore, the effects of FA on SSC-mediated bone regeneration in an irradiated bone defect animal model were explored. FA-treated irradiated SSCs or irradiated SSCs were combined with a microniche to develop an injectable SSC-microcryogel (Figure 5A).

**Figure 5.**
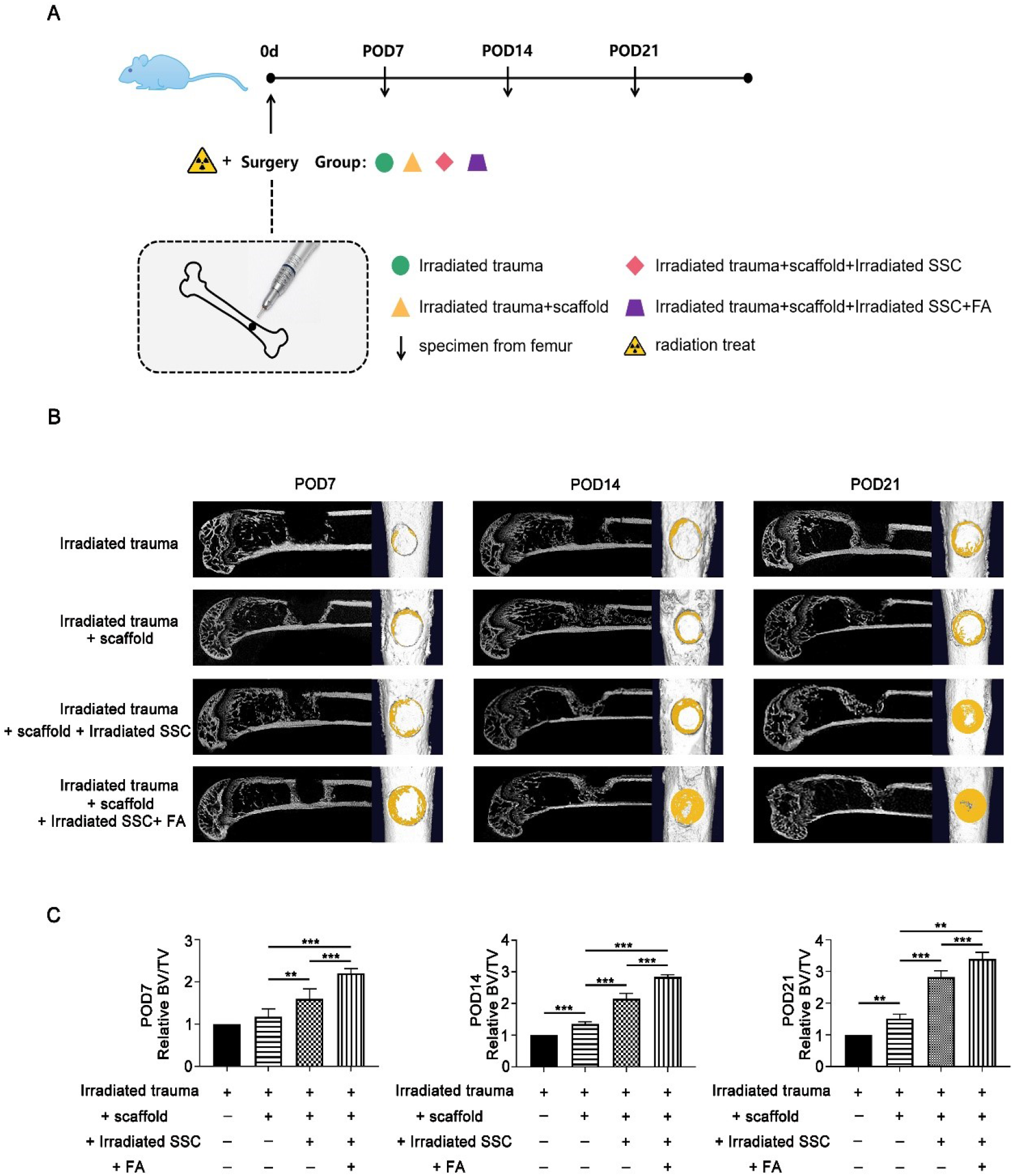
The image analysis showed that FA primed SSCs combined with microcryogel promotes bone repair in a murine irradiated bone defect model. FA-treated irradiated SSCs or irradiated SSCs were combined with a microniche to develop an injectable SSC-microcryogel (Figure 5A). As shown in Figure 5B, more newborn bone was observed in the irradiated-SSC-combined-with-microcryogel group than in the negative control group and the microcryogel group by micro-CT screening at 1, 2, and 3 weeks after SSC microcryogel transplantation. Transplantation of FA-treated irradiated SSCs combined with microcryogel yielded strengthened bone formation in the defect compared with its counterpart without FA treatment (Figure 5B). In addition, BV/TV was higher in the irradiated SSC-microcryogel group than in the negative control group and microcryogel group. The FA-treated irradiated SSC microcryogel group showed a greater increase in BV/TV than the irradiated SSC-microcryogel group (Figure 5C) (** *p* < 0.01, *** *p* < 0.001).

Bone regeneration in the defect was evaluated by micro-CT at 1, 2, and 3 weeks after SSC microcryogel transplantation. As shown in Figure 5B, more regenerated bone was observed in the irradiated-SSC-combined-with-microcryogel group than in the negative control group and the microcryogel group. Promisingly, transplantation of FA-treated irradiated SSCs combined with microcryogel yielded strengthened bone formation in the defect compared with its counterpart without FA treatment (Figure 5B). Additionally, BV/TV was higher in the irradiated SSC-microcryogel group than in the negative control group and microcryogel group. Similarly, the FA-treated irradiated SSC microcryogel group showed a greater increase in BV/TV than the irradiated SSC-microcryogel group (Figure 5C) (** *p* < 0.01, *** *p* < 0.001). To further explore bone regeneration in the defect, HE and Masson staining as well as immunohistochemical analyses of Col-I and OCN were performed. The histological data showed that only fibrous-like connective tissues filled the defect even at week 3 engrafted with microcryogel. Small bone nodules were observed in the bone defect transplanted with the irradiated SSC microcryogel (Figure 6A). Notably, bone-like tissues filled most of the defects engrafted with FA-treated irradiated SSC microcryogels. In addition, the results of Masson’s trichrome and Col-I staining showed that FA promoted the formation of collagen fibers in new bones, which contributed to bone regeneration in defects (Figure 6B). The quantitative analysis of new bone was further performed according to the results of HE and Masson’s trichrome staining. The femur defect repair percentage was calculated by Image-Pro Plus. Figure 6C shows that new bone formation in the defect engrafted by FA-treated irradiated SSC microcryogels (28.3±1.1% at day 7, 32.6±1.9% at day 14, 38.6±2.1% at day 21) was higher than in defect grafts engrafted by irradiated SSC microcryogels (21.6±2.1% at day 7, 24.0±2.9% at day 14, 28.9±3.2% at day 21) (*, *P*<0.05, **, *P*<0.01, ***, *P*<0.001).

**Figure 6.**
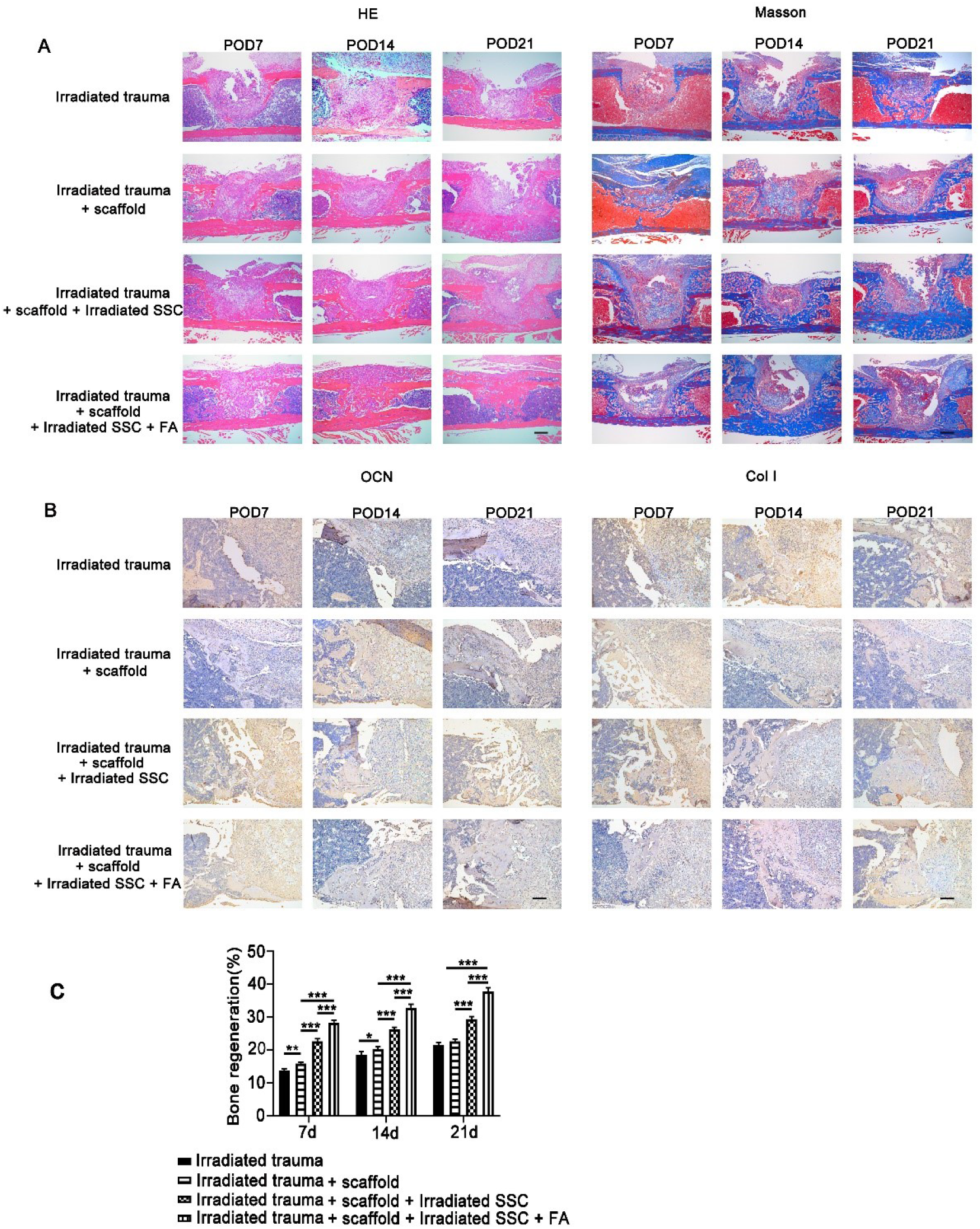
The pathological analysis of FA primed SSCs combined with microcryogel promotes bone repair in a murine irradiated bone defect model. The histological data showed that only fibrous-like connective tissues filled the defect even at week 3 engrafted with microcryogel. In addition, small bone nodules were observed in the bone defect transplanted with the irradiated SSC microcryogel (Figure 6A and 6B). Notably, bone-like tissues filled most of the defects engrafted with FA-treated irradiated SSC microcryogels. The results of Masson’s trichrome, OCN, and Col-I staining further validated that FA promoted bone regeneration in defects (Figure 6A and 6B). The quantitative analysis of new bone was further performed according to the results of HE and Masson’s trichrome staining. The femur defect repair percentage was calculated by Image-Pro Plus. Figure 6C shows that new bone formation in the defect engrafted by FA-treated irradiated SSC microcryogels (28.3±1.1% at day 7, 32.6±1.9% at day 14, 38.6±2.1% at day 21) was higher than in defect grafts engrafted by irradiated SSC microcryogels (21.6±2.1% at day 7, 24.0±2.9% at day 14, 28.9±3.2% at day 21) Bars in Figure 6A and 6B represent 200μm. *, *P*<0.05, **, *P*<0.01, ***, *P*<0.001.

### FA partially rescued the bone regenerative capacity of irradiated SSCs *in vivo* via the p38/MAPK and ERK/MAPK pathways

To explore the underlying mechanisms that control FA-SSC-mediated bone regeneration *in vivo*, specific chemical inhibitors of p38/MAPK (SB203580) and ERK/MAPK (PD98059) were added to the FA-buffered irradiated SSC culture system for 12 hours before preparing FA-treated irradiated SSC microcryogels. The bone regeneration of SSCs in each group was then evaluated. The imaging results showed that inactivation of p38/MAPK and ERK/MAPK impaired FA/SSC-mediated structural reconstitution in bone defects (Figure S3A, S3B, and S3C). Further pathological data demonstrated that SB203580 and PD98059 partially abolished the promotive effects of FA on the bone regenerative cap of irradiated SSCs *in vivo* (Figure S3A, S3B, and S3C). The data of newborn bones showed that blockage of p38/MAPK and ERK/MAPK pathways by specific inhibitors in FA treated irradiated-SSC significant abolished the bone regenerative activities in bone defects (33.5±3.6% vs 19.8±1.1% of SB203580, vs 22.1±0.9% of PD98059) (Figure S3D) (***, *P*<0.001).

## 4. Discussion

In the current study, we found that irradiation injured SSC stemness by targeting MAPK signaling *in vitro* and *in vivo*. In addition, FA was capable of maintaining SSC stemness after irradiation. Promisingly, our findings showed that FA restored the bone regenerative capacity of SSCs by activating MAPK pathways.

SSCs have been recently characterized as innate stem cells residing in skeletons that contribute to bone development, bone fracture repair, and bone remodeling^9-16,24^. However, the response of SSCs to irradiation, the role of SSCs in the settlement of radiation-induced bone injury, and the underlying mechanisms have remained largely unknown. Green et al. found that irradiation devastated the adult stem cell pools in mice preceding the collapse of trabecular bone quality and quantity^30^. Although the impact of irradiation on hematopoietic stem cells was observed, few phenotypes of skeletal tissue-derived stem cells were reported in their study. Additionally, Marecic et al. harvested dissociated progenitor cells, fracture-induced bone, cartilage, and stromal progenitor (f-BCSP) from complete femurs by collagenase digestion after bone fracture and irradiation^31^. They demonstrated that f-BCSP expansion was reduced significantly in irradiated versus nonirradiated calluses in a murine bone fracture model. Both callus development and BCSP expansion remained impaired 3 months post-irradiation^31^. Furthermore, Chandra et al. labeled all mesenchymal lineage cells in the endosteal bone marrow using a lineage tracing approach and found that radiation shifted the differentiation fate of mesenchymal progenitors to adipocytes and arrested their proliferation ability^32^. However, the stem/progenitor cells described in these reports are mainly from bone morrow or mixed with bone marrow stem cells. In our study, the bone marrow cells were removed by cavity flushing and collagenase digestion. Only SSCs, innate stem cells in bone tissues, were isolated and used in the subsequent experiments. Notably, our data showed that SSC is very sensitive to irradiation and exhibits impaired stemness, which adds novel information to understanding bone osteoporosis and hindered bone repair post-irradiation. Most importantly, the data of cellular and animal experiments suggest that SSCs may be a potential target for the settlement of irradiated bone defects.

Stem cells have been proven to be one of the important targets of drug-based therapy. Mukherjee et al. targeted the stem/progenitor population by using bortezomib, a clinically available proteasome inhibitor used in the treatment of multiple myeloma^33^. They found that bortezomib induced MSCs to preferentially undergo osteoblastic differentiation *in vitro*, and mice implanted with MSCs showed increased ectopic ossicle and bone formation when recipients received low doses of bortezomib^33^. Another study demonstrated that transplantation of pioglitazone-pretreated MSCs significantly enhanced cardiac transdifferentiation *in vitro* and improved cardiac function in a myocardial infarction model *in vivo*^34^. Additionally, treatment of MSCs with angiotensin receptor blockers has been proven to significantly improve the efficiency of cardiomyogenic transdifferentiation and cardiac function via angiogenesis^35^. Based on our understanding of SSC biology and previous research findings, we targeted irradiated SSCs *in vitro* by FA and subsequently evaluated their contribution to bone regeneration *in vivo*. First, FA was used in the current study because of its biodirectional effects of radiation protection and osteogenic promotion^26,27^. In addition, the sodium salt of FA, sodium ferulate, is a compound widely used in traditional Chinese medicine and has been approved by the National Medical Products Administration (NMPA) for clinical applications^25^. Notably, mechanistic studies have demonstrated a versatile genetic control system in mammalian cells and mice that is responsive to clinically licensed sodium ferulation^25^. Secondly, in the current study, mirocryogels were used to build stem cell-scaffold constructs that were transplanted into bone defects because the matrix is indispensable for tissue regeneration. Microcryogels have been successfully used in previous studies by virtue of their properties of cell protection, cell retention and survival, which ultimately improves the therapeutic functions of stem cells^28,29^. Thirdly, the irradiated bone defects were repaired using FA-primed allogenic SSCs and scaffolds, instead of intravenously administered FA, to recruit host SSCs because the limited endogenous SSC number in the local site may limit the efficiency of bone repair. However, FA infusion may be suitable for alleviating total body radiation-induced SSC injury and subsequent osteoporosis.

In addition to the results of cellular and animal experiments, further mechanistic data revealed the signaling pathways by which FA controls the regenerative effects of SSC. Accumulated evidence has proven that the p38 MAPK pathway is required for normal skeletogenesis in mice. Greenblatt et al. reported that mice with a deletion of the MAPK pathway members displayed remarkably reduced bone mass due to defective osteogenic differentiation. These findings suggest that p38/MAPK signaling is essential for bone formation *in vivo*^36^. In addition, ERK and p38 MAPKs cause phosphorylation of RUNX2, which promotes osteoblast differentiation. ERK also activates RSK2, which in turn phosphorylates ATF4, to enhance late-stage osteoblast synthetic functions^37^. In addition to normal skeletogenesis, MAPK signaling plays a pivotal role in bone repair. Acting as signal transductors from multiple growth factors or adhesion molecules, the MAPK pathway plays crucial functions during bone healing post-fracture^38^. Moreover, our previous work showed that inhibition of the p38/MAPK pathway caused serious osteogenic suppression^39^. Remarkably, in the present study, FA reversed radiation-induced MAPK suppression and SSC stemness injury. Further inhibition of the MAPK pathway by a specific chemical inhibitor significantly abolished the rescuing effect of FA on irradiated SSCs. Our data are consistent with previous reports that MAPK signaling plays an important role in bone regeneration and reveals the novel role of MAPK in FA-mediated anti-radiation and bone repair.

Nevertheless, we must acknowledge that there are several limitations in our study. First, although we have identified that MAPK pathways mainly contribute to the regulatory effects of FA on SSC, the potential role of other pathways, such as the TGF-β/SMAD and NF-κB pathways, should be investigated in future studies. Secondly, the contribution of endogenous SSCs and other stem cells in the host needs to be clarified, and more animal models, including genetically modified mice, may be helpful in addressing this issue. Thirdly, to explore the translational value of our findings, more drugs should be introduced into future studies, and their effects on SSCs need to be compared with FA.

## 5. Summary

In the present study, we report a new role for FA in maintaining SSC stemness after irradiation. In addition, our study reveals the underlying cellular and molecular mechanisms by which FA controls bone reconstitution post-irradiation by SSCs. Our data suggest that targeting SSCs may be a novel strategy in treating irradiated bone injury.

## Acknowledgments

This study was supported by the National Natural Sciences Grants China (No. 81871771, 81572159,81500083,) and the Beijing Natural Sciences Foundation (No. 7182123,7192203).

## Data Availability Statement

The data that support the findings of this study are available from the corresponding author upon reasonable request.

---

**Figure S1 The influence of irradiation on adipogenic differentiation of SSCs**

No obvious influences of irradiation (2Gy) on adipogenic droplet formation (Figure S1A) and gene expression of adipogenic differentiation of SSC (Figure S1B) were observed in SSCs.

Bars in Figure S1A represent 200μm.

**Figure S2 The general change of irradiation-induced gene expression in SSCs**

The data in Figure S2A and S2B present the differentially expressed genes in heatmap and volcano plot format between SSC and irradiated SSC. The results of mRNA sequencing showed that a total of 638 genes were upregulated, and 198 genes were downregulated in the irradiated SSC group compared with the SSC control group (2 Gy, n = 3).

**Figure S3 Blockage of p38/MAPK and ERK/MAPK in SSCs partially abolished FA primed bone regeneration in irradiated bone defects**

The imaging results of the bone regeneration of SSCs in each group showed that inactivation of p38/MAPK and ERK/MAPK impaired FA/SSC-mediated structural reconstitution in bone defects (Figure S3A, S3B, and S3C). Further pathological data demonstrated that SB203580 and PD98059 partially abolished the promotive effects of FA on the bone regenerative cap of irradiated SSCs *in vivo* (Figure S3A, S3B, and S3C). Moreover, blockage of p38/MAPK and ERK/MAPK pathways by specific inhibitors in FA treated irradiated-SSC significant abolished the bone regenerative activities in bone defects (33.5±3.6% vs 19.8±1.1% of SB203580, vs 22.1±0.9% of PD98059) (Figure S3D). Bars in Figure S3C represent 200μm. *** *p* < 0.001.

